# Role of Ribeye PXDLS/T-binding cleft in normal synaptic ribbon function

**DOI:** 10.1101/2023.12.12.571266

**Authors:** Jie Zhu, Caixia Lv, Diane Henry, Stephen Viviano, Joseph Santos-Sacchi, Gary Matthews, David Zenisek

## Abstract

Non-spiking sensory hair cells of the auditory and vestibular systems encode a dynamic range of graded signals with high fidelity by vesicle exocytosis at ribbon synapses. Ribeye, the most abundant protein in the synaptic ribbon, is composed of a unique A domain specific for ribbons and a B-domain nearly identical to the transcriptional corepressor CtBP2. CTBP2 and the B-domain of Ribeye contain a surface cleft that binds to proteins harboring a PXDLS/T peptide motif. Little is known about the importance of this binding site in synaptic function. Piccolo has a well-conserved PVDLT motif and we find that overexpressed Ribeye exhibits striking co-localization with Piccolo in INS-cells, while two separate mutants containing mutations in PXDLS/T-binding region, fail to co-localize with Piccolo. Similarly, co-transfected Ribeye and a piccolo fragment containing the PVDLT region co-localize in HEK cells. Expression of wild-type Ribeye-YFP in zebrafish neuromast hair cells returns electron densities to ribbon structures and mostly rescued normal synaptic transmission and morphological phenotypes in a mutant zebrafish lacking most Ribeye. By contrast, Ribeye-YFP harboring a mutation in the PXDLS/T-binding cleft resulted in ectopic electron dense aggregates that did not collect vesicles and the persistence of ribbons lacking electron densities. Furthermore, overexpression failed to return capacitance responses to normal levels. These results point toward a role for the PXDLS/T-binding cleft in the recruitment of Ribeye to ribbons and in normal synaptic function.

**Significance statement:** Hair cell synaptic ribbons are evolutionarily conserved structures that appear dense in electron micrographs, extend from the release site into the cytoplasm and tether synaptic vesicles for release. Here, we show that Ribeye, the major component of the synaptic ribbon electron density is directed to ribbons via its PXDLS/T-binding site and that mutant isoforms of Ribeye with point mutations disrupting these interactions fail to rescue normal function in mutant zebrafish lacking most Ribeye.

## Introduction

Sensory neurons of the auditory, vestibular and visual system utilize ribbon synapses to transmit signals at high spatial and temporal resolution. The synaptic ribbon is a protein-based structure in which the most-abundant protein is Ribeye (Schmitz et al., 2000; Zenisek et al., 2004; Kantardzhieva et al., 2011). The functional role of Ribeye in ribbon synapses remains unclear. Work in mouse retinal ribbons (Maxeiner et al., 2016; Okawa et al., 2019; Grabner and Moser, 2021; Mesnard et al., 2022) and hair cells in zebrafish (Sheets et al., 2011; Lv et al., 2016) and mouse (Becker et al., 2018; Jean et al., 2018) have revealed that Ribeye is required for normal ribbon morphology, presynaptic protein localization and normal vision and hearing, but effects on synaptic transmission have varied.

Ribeye arises from an alternative start site to the *CtBP2* gene. Ribeye contains an A domain that is unique to Ribeye and a highly conserved B domain, which is identical to CtBP2, except for the 20 N-terminal amino acid that include a nuclear localization sequence (Schmitz et al., 2000). CtBP2 functions as a transcriptional co-repressor that regulates transcription in a redox-dependent manner in vertebrates (Zhang et al., 2000).

CtBP2 and the closely related CtBP1 have been studied in great detail (reviewed in (Chinnadurai, 2002; Stankiewicz et al., 2014; Blevins et al., 2017) and the previous work can serve as a guide for generation of hypotheses on the function of the Ribeye B-domain. The CtBPs do not interact directly with DNA, but instead exert their transcriptional activity through interactions with other proteins. Most of the known interactions of the CtBP protein are with proteins containing variations of a PXDLS/T motif (Chinnadurai, 2002). These PXDLS-motif proteins interact with CtBP by binding to a surface cleft in the CtBP proteins (Zhang et al., 2000; Chinnadurai, 2002; Nardini et al., 2003) and mutations in the PXDLS interactions site on CtBPs disrupt CtBP co-repressor activity.

Despite the importance of these interactions to CtBP function, it remains unknown whether similar interactions are important to Ribeye function. Of note, the only ribbon protein known to harbor a PXDLS/T motif is Piccolino, a ribbon specific isoform of the active zone protein Piccolo (Regus-Leidig et al., 2013; Regus-Leidig et al., 2014; Muller et al., 2019). In this study we find evidence for an interaction between zebrafish Ribeye and Piccolo/Piccolino, consistent with previous work with mammalian Ribeye and Piccolino (Muller et al., 2019).

To further probe the importance of the PXDLS/T binding cleft in ribbon synapse function, we took advantage of a line of zebrafish that expresses little Ribeye in the hair cells in lateral line neuromasts(Lv et al., 2016). These fish exhibit ‘ribbon ghosts’ in neuromasts that lack the electron dense core and poorly localize to synapses but maintain normal numbers of vesicles. Membrane capacitance measurements reveal that they have slightly elevated exocytosis and calcium current. We find here that overexpression of Ribeye containing a mutation to the PXDLS-binding cleft in this background fails to rescue either the morphological or electrophysiological phenotype. Our results indicate that an intact PXDLS-binding cleft is necessary for normal hair cell synaptic ribbon function.

## Materials and Methods

### Experimental Design

#### Ribeye cloning and plasmid constructions

Ribeye mRNA was purified from the retinas of zebrafish. Oligo-dT primed cDNA was synthesized using Superscript II Reverse Transcriptase (Invitrogen). 5’ and 3’ oligonucleotides were designed based on homology of predicted zebrafish sequence (from assembled genomic sequence) according to the sequence from mice. Products were isolated by agarose gel electrophoresis, and sent to sequence. A sequence was found to be identical to published *ribeye as* (accession AY878350.1) sequence previously published (Wan et al., 2005). The products was sub-cloned into pCDNA3 (Invitrogen) for cell culture. Full length Rattus Ribeye cDNA was subcloned in to pEGFP-N1 vector through NheI and Age I. Various PXDLS/T-binding motif base mutations were introduced using a commercial site-directed mutagenesis kit (QuikChange; Stratagene). Chimeric gene constructs were constructed by PCR-based approaches with point mutation primers and GFP tag was fused on the C-terminal to make rat Ribeye -EGFP (rRE-EGFP). To further study the PXDLS/T-binding motif in rat Ribeye, we then made point mutations in the Ribeye PXDLS/T-binding motif and generated two mutations. One changes the 583 Aspartic acid into Alanine (D583-6A-GFP) and another changes 615 Valine into Arginine(V615R-GFP). To study the interactions of Ribeye and Piccolo (2622-2937) in cell lines, rattus Piccolo (2622-2937) DNA was cloned into vector pCDNA3.1-myc-his(C) vector (Invitrogen).

To study the PXDLS/T binding-motif of Ribeye function in synapses, we then sub-cloned the zebrafish *ribeye as and mutated ribeye PXDLS/T-binding mutants* were concatenated to YFP by PCR-based approach and then sub-cloned into a myo6b promoter vector (obtained from Dr. Lavinia Sheets, Washington University and Dr. Katie Kindt, NIDCD, National Institute of Health) in the Tol 2 system (Suster et al., 2009). This encodes myo6b promoter specific for zebrafish hair cells and a myosin light chain 2 promoter that drives expression of GFP in the heart. We co-injected one-cell embryos with plasmid DNA (50ng/μl) described with mRNA encoding tol2 transposase (60ng/μl) to make mosaic zebrafish.

Standard molecular methods were used for manipulation of plasmid DNA from E. coli. Point mutation were made in *ribeye a* (1113g1114t to 1113c1114g), resulted in a coding Valine into arginine (V338R) in Ribeye PXDLS motif, generated a mutated Ribeye.

To make rRE-E1AC, we concatenated the C terminal 63 amino acids (after changing the nuclear localization signal, KRPRP to AAPAP) of E1A (a kind gift from Daniel DiMaio, Yale University) to the C-terminal of Ribeye. HA tagged (YPYDVPDYA) Ribeye (rRE-HA) was made as a control.

#### Peptides synthesis

Ribeye-binding peptides: rhodamine Ribeye binding peptides (rhod-EQTVPVDLSVARPR-cooh) (Zenisek et al., 2004; Francis et al., 2011) or fluorescein Piccolo-peptides (fluor-IEDEEKPVDLTAGRRA) were synthesized and purified (>90% purity) by GenScript. Peptides were solubilized and frozen as a stock solution of 20 mM in water and diluted to a final concentration of 10 µM.

#### Bipolar cell dissociation

Cells were prepared as previously described (Coggins et al., 2007) in accordance with IACUC at Yale University. Following enucleation, corneas were removed and eye cups placed in solution containing 20 units/mL hyluranidase 0.5 Ca^2+^, 120 NaCl, 2.5 KCl, 1.0 MgCl_2_, 0.5 CaCl_2_, 10 Glucose, 10 HEPES adjusted to a pH = 7.4. Next the neural retina was removed from the eyecup and digested for 35 min at room temperature in papain (35 u/mL; Sigma) containing cysteine (0.5 mg/mL) and 0.5 Ca^2+^, 120 NaCl, 2.5 KCl, 1.0 MgCl_2_, 0.5 CaCl_2_, 10 Glucose, 10 HEPES adjusted to a pH = 7.4. The digested material was rinsed several times and stored in a 12 − 14°C, oxygenated refrigerator. Cells were dissociated by trituration on to a high refractive index coverslip using a fire polished Pasteur pipette. Fluorescent peptides were introduced via whole-cell patch pipette containing (in mM): 120 Cesium Gluconate, 4.0 MgCl_2_, 10 HEPES, 10 TEA-Cl, 0.5 EGTA, 0.5 Na_2_GTP, 4.0 Na_2_ATP, adjusted to pH 7.3 with CsOH. Glass recording electrodes were prepared from thick-walled pipettes that were pulled to a resistance of 8-12 MΩ.

#### Cell culture

PC-12 and Ins-1 cells were transfected using lipofectamine 2000 as per manufacturers’ instructions (Invitrogen, Carlsbad, California, USA). After transfection, cells were allowed to recover in medium for 2 hrs before being harvested using Trypsin/EDTA and replated on coverslips at a density of 1.5 × 10^5^ cells cm^−2^. Cells were allowed to adhere and spread overnight before imaging.

#### Mosaic transgenic zebrafish

Zebrafish were kept in accordance with the Yale University Animal Care and Use Committee guidelines. Mutant alleles used in this study are the *ribeye [a(*Δ*10)/b(*Δ*7)]* double-homozygous mutant zebrafish, which have been described previously (Lv et al., 2016). To generate mosaic transgenic fish, plasmid DNA (150 ng/μl) and RNA encoding tol2 transposase (60 ng/μl) (Kawakami 2004; Suster et al. 2009) were injected into one-cell-stage embryos. Mosaic transgenic animals were screened as green hearts by fluorescence microscopy 2-3 days post-fertilization.

#### Immunohistochemistry

Zebrafish larvae from 4 to 6 dpf (day after post fertilization) were anaesthetized and fixed in 4% paraformaldehyde at 4 ⁰C overnight. Fish larvae were washed by PBS three times then permeabilized in acetone for 7 minutes at −20°C, and blocked in 1% bovine serum albumin (v), 3% normal goat serum (v/v), and 0.1% Triton X-100 in PBS at room temperature for 2 hours, followed by overnight incubation with primary antibodies in PBST at 4°C. The second morning, the samples were washed 3 times with PBST, and incubated in diluted secondary antibody for 2 hours at room temperature. Then fish were mounted in mounting solution (ProLong Gold antifade reagent with DAPI, Invitrogen, USA).

#### Antibodies

The primary antibodies used for cell staining and zebrafish were CtBP [CtBP (B-3, Santa Cruz Biotechnology, Inc., Santa Cruz, CA, USA)], RIM2 (anti-Rim2 PDZ domain, Synaptic Systems, Goettingen, Germany), CaV1.3(L-type Ca++CP, H160, Santa Cruz Biotechnology, USA), Bassoon (ab76065, Abcam, Cambridge, MA, USA), and c-myc (9E10) (sc-40, Santa Cruz Biotechnology, Inc., Santa Cruz, CA, USA) and Piccolo (Anti-Antibody [6H9-B6] and StressMarq Biosciences INC; Anti-Piccolo antibody (ab20664) Abcam, Cambridge, MA, USA), anti-green fluorescent protein mouse IgG1(Invitrogen A11121); anti-green fluorescent protein mouse IgG2a(Invitrogen 1037264)GFP monoclonal Antibody(clontech632375) were diluted 1:200. The secondary antibodies were Alexa Fluor 568 goat anti-rat IgG(H+L) (Invitrogen Lot:430228), Alexa Fluor 568 goat anti-rat IgG(H+L) (Invitrogen A11077); Alexa Fluor 568 goat anti-rabbit IgG(H+L) (Invitrogen A11011); Alexa Fluor 488 goat anti-mouse IgG(H+L) (Invitrogen A11001) were diluted 1:200.

The primary antibodies used for zebrafish whole mount staining were CtBP (CtBP B-3, Santa Cruz Biotechnology, USA) antibody and pan-MAGUK (Neuromab) antibody were diluted 1:500. The GFP antibody (anti-GFP, rabbit lgG fraction, Invitrogen, USA) was diluted 1:1000. The secondary antibodies (Alexa Fluor 488/594 goat anti mouse/rabbit IgG(H+L), Invitrogen, USA) were diluted 1:1000. They are Alexa Fluor 488 goat anti rabbit IgG(H+L) (Invitrogen A32731), Alexa Fluor 594 goat anti rabbit IgG(H+L) (Invitrogen A11037), Alexa Fluor 488 goat anti mouse IgG(H+L) (Invitrogen A11001), Alexa Fluor 594 goat anti mouse IgG(H+L) (Invitrogen A11032).

#### Confocal Imaging

Cell culture images were taken using a Zeiss (Germany) LSM 780 laser-scanning confocal. All images for any comparisons were taken using the same settings via a Plan-Apochromat 63x/1.40 Oil DIC M27 objective. Image J was used for protein co-localization analysis, with the methods described by McMaster Biophotonics Facility (www.macphotonics.ca). To quantify the colocalization of the proteins, we use the intensity correlation quotient (ICQ: McMaster Biophotonics Facility) measurement to indicate the degree of which the intensity of two colors in an image varies in synchrony relative to the means, following the approach developed by Stanley and colleagues (Li et al., 2004). For random staining, the ICQ value is around 0; for segregated staining, 0>ICQ≥-0.5, and for dependent staining, 0<ICQ≤0.5. ICQ measurements vary from −0.5 to 0.5, where positive values indicate dependent staining with higher numbers indicating a greater degree of synchrony between the two colors.

Whole-mounted zebrafish lateral lines were imaged with Zeiss (Germany) LSM 800 laser-scanning confocal microscope using Plan-Apochromat 63x/1.40 Oil DIC M27 objective and 1x or 5x digital zoom. GFP samples were imaged using a 488 nm laser for excitation and 495-555nm emission. Alex-594 samples were images using a 561nm excitation and 580-680nm emission, with best signal scanning in the software set up. Images of 1024×1024 pixels were acquired at a plane scanning mode with 4 line averages. Images were stored as 8-bit RGB tiff files for further analysis and processing.

#### Zebrafish in-vivo hair cell patch-clamping

Hair cell recordings were performed as previously described (Ricci et al., 2013; Lv et al., 2016). Briefly, zebrafish larvae dpf 4-9 were anesthetized in Tricane(168mg/l) and paralyzed in Tubocurarine(10 μM), mounted in a recording chamber and tied down using dental floss. Recordings were made under an upright Olympus microscopy with an Axon 200B amplifier,an Axon DD1322 digitizer. All recordings were made with jClamp software. To access hair cells for whole-cell recordings, supporting cells surrounding the hair cells were first removed and cleaned by a large bore pipette (3-5 MΩ). All recordings were made with jClamp software (Scisoft).

Cells were held at a membrane potential of −80 mV. Extracellular solution was as follows (in mM): 125 NaCl, 1.0 KCl, 2.2 MgCl_2_, 3 CaCl_2_, 10 HEPES, 6 D-glucose, 285 mOsm, pH 7.4. Pipette solution was (in mM): 90 CsCl, 20 TEA, 5 Na_2_ATP, 3.5 MgCl_2_, 10 HEPES, 1 EGTA, 260 mOsm, pH 7.3. Pipette resistance was typically 7-10 MΩ. Capacitance measures were made using the dual sine admittance technique (Santos-Sacchi, 2004). Capacitance data were smoothed by averaging 10 adjacent points in time before averaging across cells.

#### Electron Microscopy

Electron microscopy methods were adapted from previously published protocols (Obholzer et al. 2008)(Lv et al., 2016). Briefly, 4-8 dpf zebrafish larvae were fixed overnight in 3% glutaraldehyde and 1.5% paraformaldehyde in 0.1M phosphate buffer overnight at 4°C. After rinsing in 0.1M PBS, samples were stained with 1% osmium, dehydrated in serial washes of 50%, 80%, 95% and then 100% ethanol. Fish were embedded in EMbed-812 (EMS). Hair cells from anterial neuromasts were imaged on Tecnai Biotwin electron microscope.

To identify hair cells overexpressing transgenes, only fish exhibiting the most wide-spread expression of GFP in neuromasts were chosen for electron microscopy analysis. This approach was validated as effective, since we were reliably able to identify densities in Ribeye over-expressing *ribeye [a(*Δ*10)/b(*Δ*7)]* double homozygous fish, but never in *ribeye [a(*Δ*10)/b(*Δ*7)]* where transgenes were not being overexpressed.

#### Statistical analysis

All the data are expressed as averages ± s.e.m. unless otherwise noted. Two sample Student’s t-test were used to test for statistical significance in pair-wise comparisons. ANOVAs were used when more than one condition was compared.

## Results

### RIBEYE expressions in Ins-1 cells

As a starting point for this study, we made the observation that Ribeye-EGFP overexpressed in Ins-1 cells exhibit small and more uniform spots of 0.13±0.01 µm^2^; n = 28 (Figure 1) that tended to aggregate near the membrane, while the expression pattern in HEK-293 cells often showed large aggregates within and throughout the cytoplasm of 0.27±0.01µm^2^; n = 19 showed larger puncta and may flow inside the cytoplasm Because Ins-1 cells are derived from pancreatic beta cells, which secrete both neurotransmitters and insulin, we hypothesized that INS-1 cells may harbor proteins that interact with Ribeye affecting the shape and size of the aggregates. Pancreatic beta cells express proteins such as Piccolo, RIM2, Bassoon, and the L-type calcium channel CaV1.3 (Liu et al., 2003; Ohara-Imaizumi et al., 2005; Jacobo et al., 2009).

**Figure 1.**
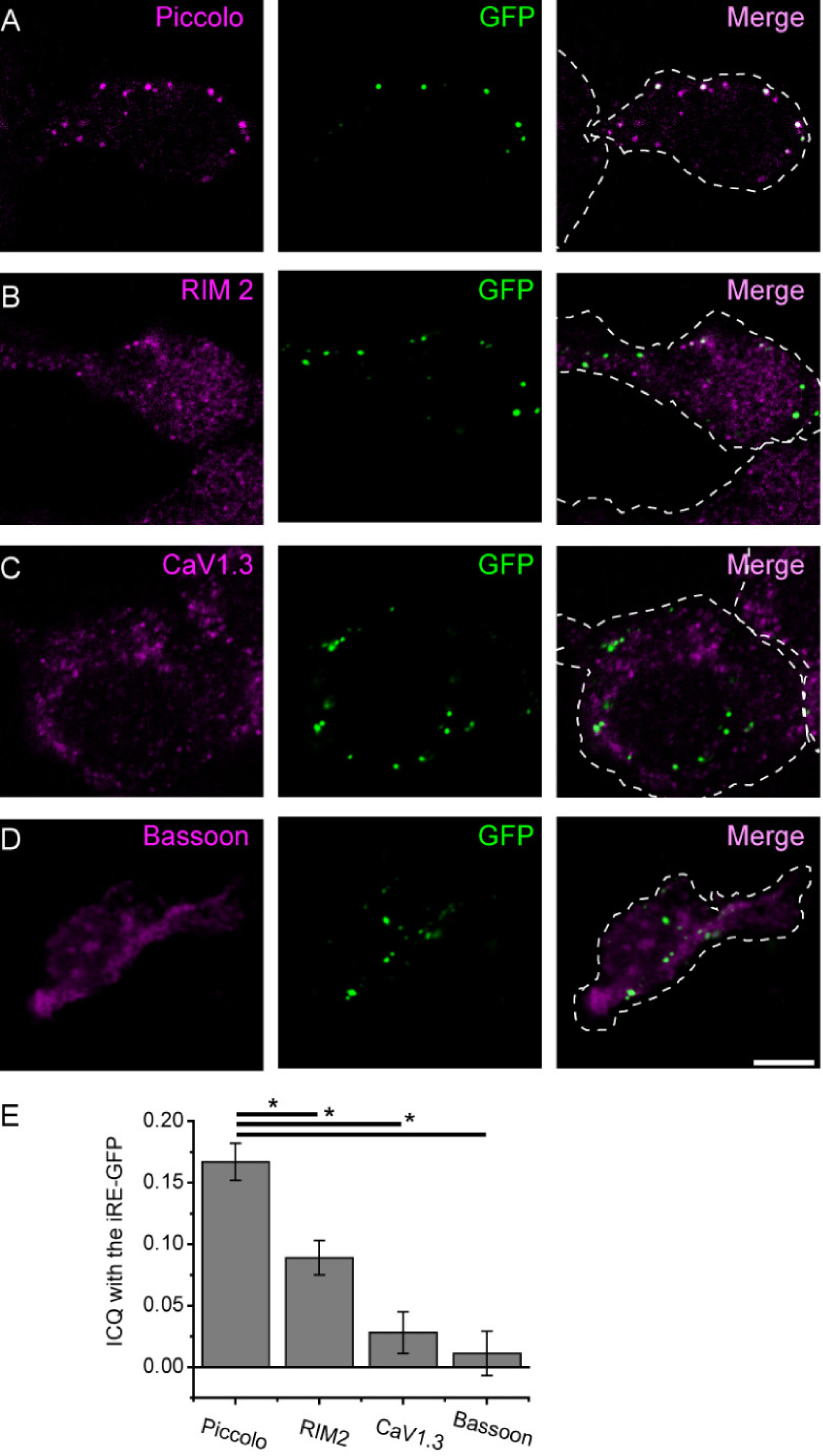
Colocalization of overexpressed Ribeye with presynaptic proteins in INS-1 cells. INS-1 cells were transfected with rat Ribeye-GFP and then immuno-stained for Ribeye and several presynaptic proteins. (A-D) show example images from co-staining for Ribeye and Piccolo (A), RIM2 (B), CaV1.3 (C) and Bassoon (D). For each set of images, magenta denotes the presynaptic protein of interest, green indicates GFP and the right panel merges both colors. Note the striking degree of colocalization between spots of Piccolo and GFP in A; scale bar: 5µm applies to all images. (E) Colocalization was quantified by determining the ICQ value. The ICQ varies from −0.5 to 0.5, with the higher numbers indicating better correlation between the two channels. For each protein pair, an ICQ value was determined for Piccolo n=17, RIM2 n=14, CaV1.3 n=4 and Bassoon n=6 transfected cells images and averaged. *P<0.05.

To investigate which proteins may colocalize and possibly interact with Ribeye, we performed immuno-staining with several ribbon-associated proteins in INS-1 cells. Immunocytochemistry revealed that Ribeye co-localizes with Piccolo, a cytomatrix protein at synaptic ribbons (Dick et al., 2001; Limbach et al., 2011). Similarly, rRE-EGFP showed partial colocalization with RIM2, CaV1.3 and Bassoon (figure 1A-D), proteins also associated with synaptic ribbons(Brandstatter et al., 1999; Dick et al., 2003; Schnee and Ricci, 2003; tom Dieck et al., 2005; Joiner and Lee, 2015). To quantify the degree of colocalization between Ribeye and other presynaptic proteins, we calculated the intensity correlation quotient (ICQ:see methods) according to each protein (Figure 1E). ICQ measurements vary from −0.5 to 0.5, where positive values indicate dependent staining with higher numbers indicating a greater degree of synchrony between the two colors. As expected from observation of the images, we found that the ICQ for Ribeye and Piccolo (0.167±0.015, n = 17 images) was higher than the ICQ values for Ribeye and RIM2 (0.089±0.014; n = 14), Cav1.3 (0.028±0.017; n = 4) and Bassoon (0.011±0.018; n = 6). These results indicate a high degree of colocalization between Ribeye and Bassoon, a lesser degree of colocalization between RIM2 and Ribeye and no significant co-localization of Ribeye with either Cav1.3 or Bassoon.

### PXDLS/T motif of Ribeye is required for the localization with endogenous Piccolo in INS-1 cells

Ribeye is composed of a self-aggregating A domain and a B domain that is identical to CtBP2 (Schmitz et al., 2000). Previous studies showed that, CtBP1 and CtBP2, interact with most of their binding partners via a PXDLS/T motif binding cleft (Chinnadurai, 2002; Kuppuswamy et al., 2008; Zhang et al., 2002; Zhang et al., 2000; Zhao et al., 2007). Fluorescent peptides containing the PXDLS motif modeled after CtBP-binding proteins bind to the synaptic ribbon and have been used as labels for fluorescent imaging, indicating that this binding site is retained and available in Ribeye in synaptic ribbons. Notably, Piccolo contains a PVDLT sequence, which is conserved across all vertebrates and is retained within the ribbon-synapse specific isoform, known as Piccolino (Regus-Leidig et al., 2013; Regus-Leidig et al., 2014). Moreover, more recently Muller et al., showed that Ribeye can bind to a region of Piccolino that contains the PVDLT motif.

Previous studies have shown that CtBP1 mutations DGRD34-37A and V66R disrupt the PXDLS/T-binding ability (Kuppuswamy et al., 2008). Therefore, we made the corresponding mutations in rat Ribeye, D583-6A EGFP and V615R EGFP, respectively, to test whether they disrupt localization to piccolo in INS-1 cells. As with WT-Ribeye-EGFP, both D583-6A and V615R exhibited a punctate pattern, indicating that these mutations did not prevent aggregation (Figure 2 A-C left panels). However, both D583-6A and V615R failed to co-localize with Piccolo (Figure 2 BC right merge panels). ICQ value showed that compared to wild type Ribeye (ICQ=0.176±0.012, n=18), D583-6A has a value as 0.060±0.018 (n=12) and V615R has a value of 0.039±0.011 (n=20), indicating that disrupting the PXDLS/T-binding motif dramatically reduced Ribeye colocalization with Piccolo (Figure 2D).

**Figure 2.**
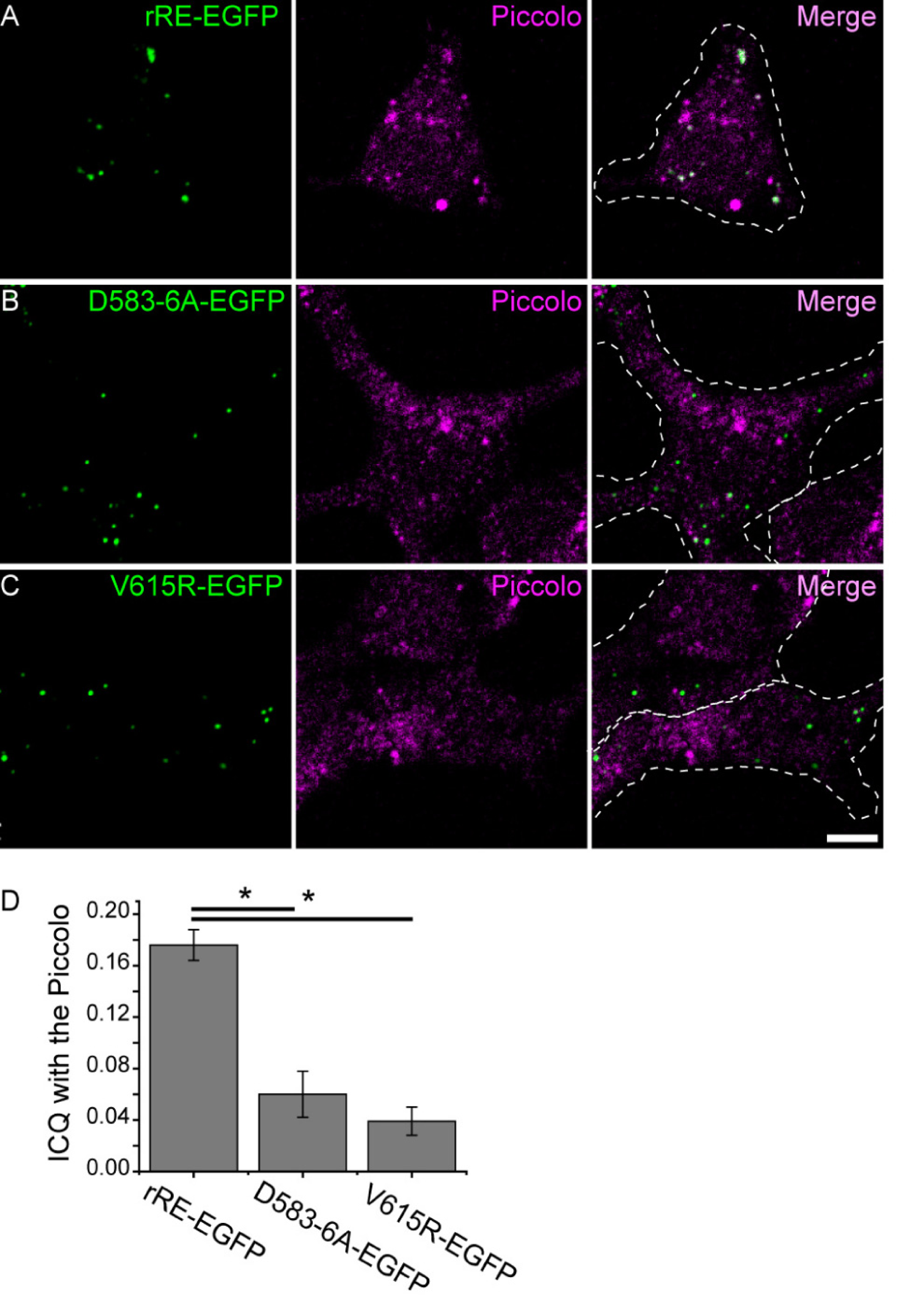
Ribeye-Piccolo colocalization requires intact PXDLS/T-binding cleft. (A-C): Examples of cells transfected with Ribeye-EGFP (A) or Ribeye-EGFP with two different mutations in the PXDLS/T binding cleft (B-C). Green denotes EGFP, whereas Magenta denotes Piccolo localization. D, Degree of correlation was calculated using the ICQ quantification. Wild type Ribeye n=18, D583-6A n=12, V615R n=20 cells. Scale bar: 5 µm. *P<0.05.

Work from the Chinnadurai laboratory found that mutations to the PXDLS/T-binding cleft may have more widespread effects on interactions beyond those directly binding to this region (Kuppuswamy et al., 2008). A “cleft filled” construct where CtBP is concatenated to the CtBP-binding motif of adenovirus E1A has been previously developed to specifically block interactions of the PXDLS/T motif binding cleft with other proteins (Kuppuswamy et al., 2008). Therefore, we took a similar approach in our experiments. We transfected cells with either rat Ribeye containing an HA tag (rRE-HA) or Ribeye with the c-terminus of E1A concatentated to the C-terminus of ribeye (rRE-E1AC) into INS-1 cells and examined the colocalization of Ribeye and Piccolo. In contrast to rRE-HA, which forms aggregates and co-localizes with Piccolo (Figure 3A), rRE-E1AC showed a much more diffuse distribution and co-localization with Piccolo was significantly reduced (Figure 3B). As expected from the images, ICQ analysis revealed a lower degree of colocalization for the cleft-filled mutant (ICQ = 0.21 ± 0.02’ n = 12) than the wild-type (ICQ = 0.44±0.02 *n* = 12) (*p* < .001). Of note, the ICQ value is significantly above 0 for the cleft-filled mutant, suggesting some degree of colocalization. These results suggest that piccolino may be able to compete with the c-terminal concatenated E1A, albeit poorly, for binding to Ribeye. This result is consistent with the 1-100 mM affinity exhibited by single linear peptides for CtBP-proteins (Molloy et al., 2000), compared with the nM affinity exhibited by full-length proteins(Molloy et al., 2001).

**Figure 3.**
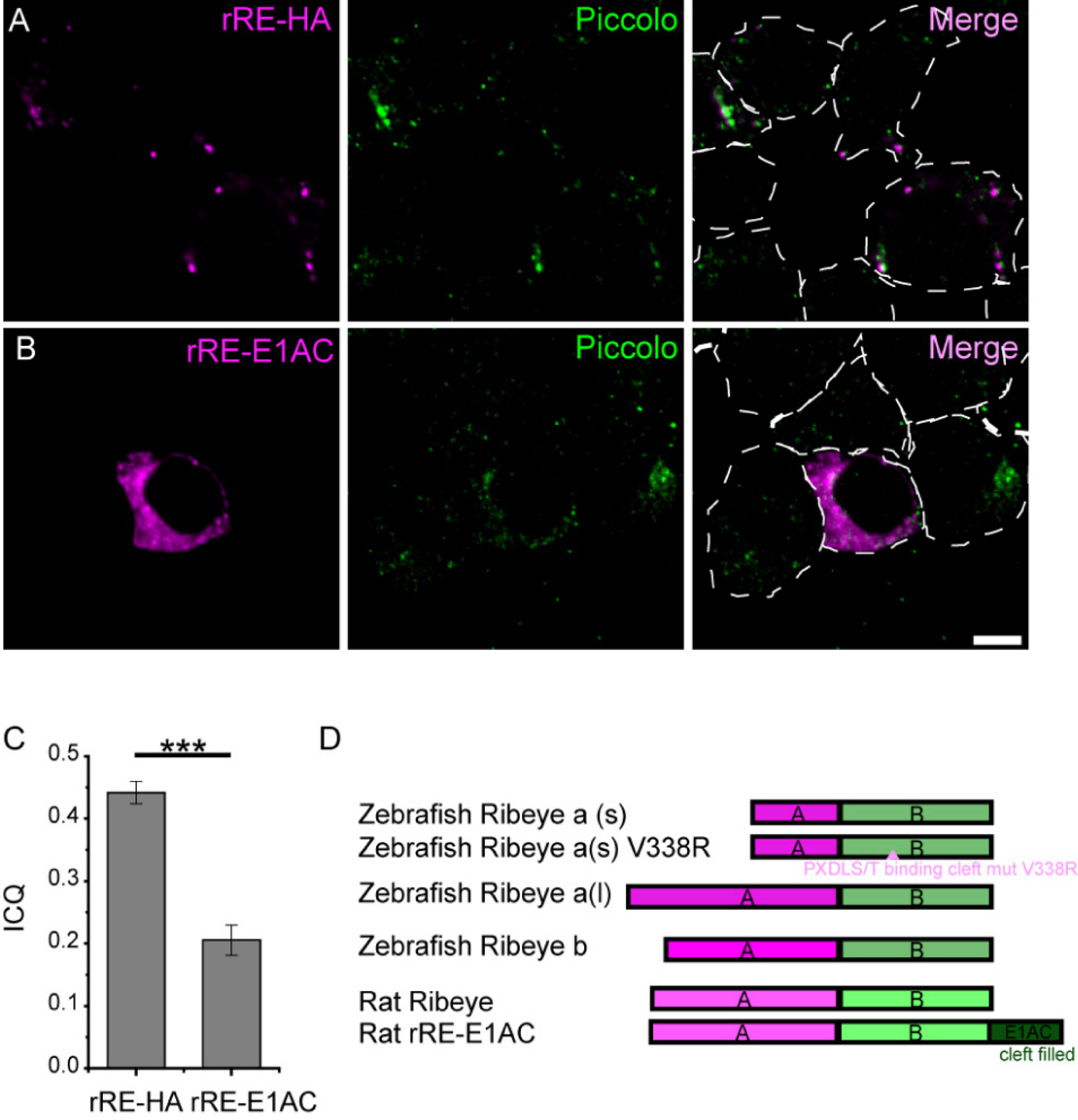
Cleft-filled Ribeye mutant fails to localize to Piccolo in INS-1 cells. INS-1 cells were transfected with construct for expression of either rat Ribeye-HA (A) or a ‘cleft-filled’ rat Ribeye (B), which has the E1A sequence concatenated to its C-terminus. E1A binds to the B-domain of Ribeye and thus is expected to compete with Piccolo for binding at the PXDLS/T binding cleft. Ribeye constructs are shown in magenta and piccolo staining is shown in green. Scale bar is 5 µm.

In summary, our results demonstrate that the PXDLS/T-binding cleft of Ribeye is critical for its interaction with Piccolo.

### Ribeye localizes to Piccolo (2622-2937)

To further investigate this interaction, next we used a fragment of Piccolo (2622-2937) that contains the PVDLT sequence to specifically test the interactions with Ribeye in HEK cells. Unlike INS-1 cells, HEK cells lack detectable expression of synaptic ribbon proteins such as RIM1, RIM2, Piccolo, bassoon or CaV1.3 (data not shown). We generated a construct containing a c-myc tag at the C-terminal of Piccolo (2622-2937) and co-transfected it with Ribeye. We found that without Ribeye, the expression of Piccolo (2622-2937) was nearly un-detectable and showed dim diffuse expression (Figure 4A). When Piccolo (2622-2937)-myc was co-transfected with Ribeye-EGFP or Ribeye-HA, the Piccolo (2622-2937)-myc fragment exhibited a punctate expression pattern that colocalized with Ribeye (Figure 4B and C) (ICQ=0.178±0.056, n=16). By contrast, anti-myc staining exhibited weak expression when co-transfected with rRE-E1AC (Figure4 D). Together, our results suggest that Ribeye interacts with Piccolo (2622-2937) fragment and disrupting the PXDLS/T motif binding cleft prevents this interaction.

**Figure 4.**
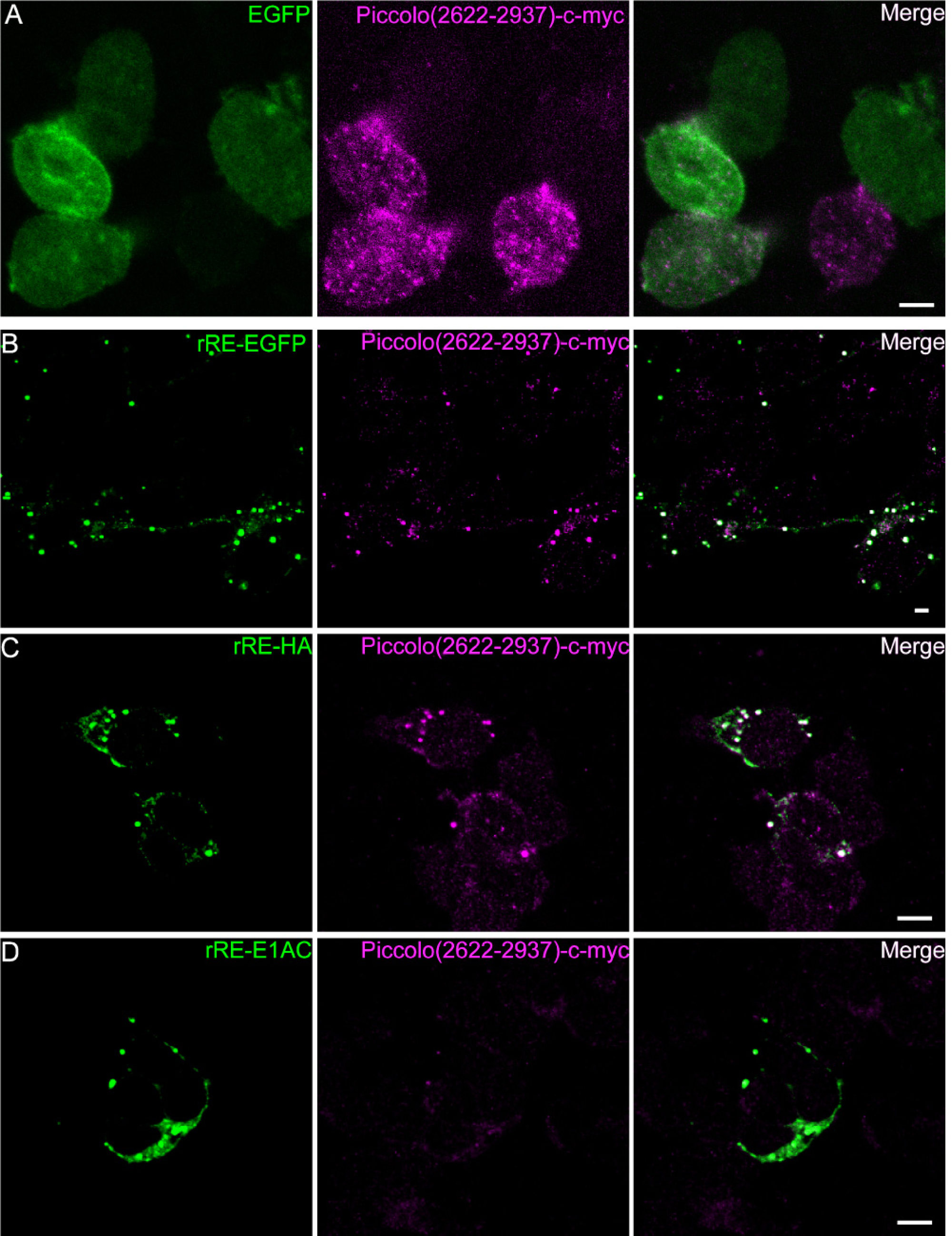
Co-expression of Ribeye and Piccolo (2622-2937) in HEK cells shows striking colocalization that is disrupted in cleft-filled mutant. HEK cells were co-transfected with a fragment of Piccolo containing the PVDLT sequence (Piccolo(2622-2937)) and either EGFP (A) or variants of Ribeye (B-D). Piccolo(2622-2937) exhibited striking colocalization with Ribeye-EGFP (B) and Ribeye-HA (C), but not the cleft filled Ribeye (D; rRE-E1AC-HA) or GFP. Scale bar: 5 µm

Previous work from our laboratory has demonstrated that fluorescently-tagged short peptides containing a PXDLS/T-motif localize to ribbons (Zenisek et al., 2004; Francis et al., 2011). These fluorescent peptides appear as prominent spots when introduced into goldfish bipolar cells (Figure 5 A) via a patch pipette and imaged using TIRF microscopy (Figure 5C). We tested whether a peptide made up of a short stretch of Piccolo sequence that includes the PVDLT motif shows the same pattern (Figure 5 D). The Piccolo derived sequence (PCLO-14) exhibited bright spots that were readily visible and co-localized with the previously verified Ribeye-binding peptide (Zenisek et al., 2004) in both TIRF in goldfish bipolar cells and confocal in mouse bipolar cells, confirming that this peptide is a good substrate for ribbon binding. (Figure 5). As expected, ICQ analysis revealed a high degree of colocalization between PCLO-14 and the Ribeye-binding peptide (ICQ = 0.46 +/− .01; n = 4 TIRF images).

**Figure 5.**
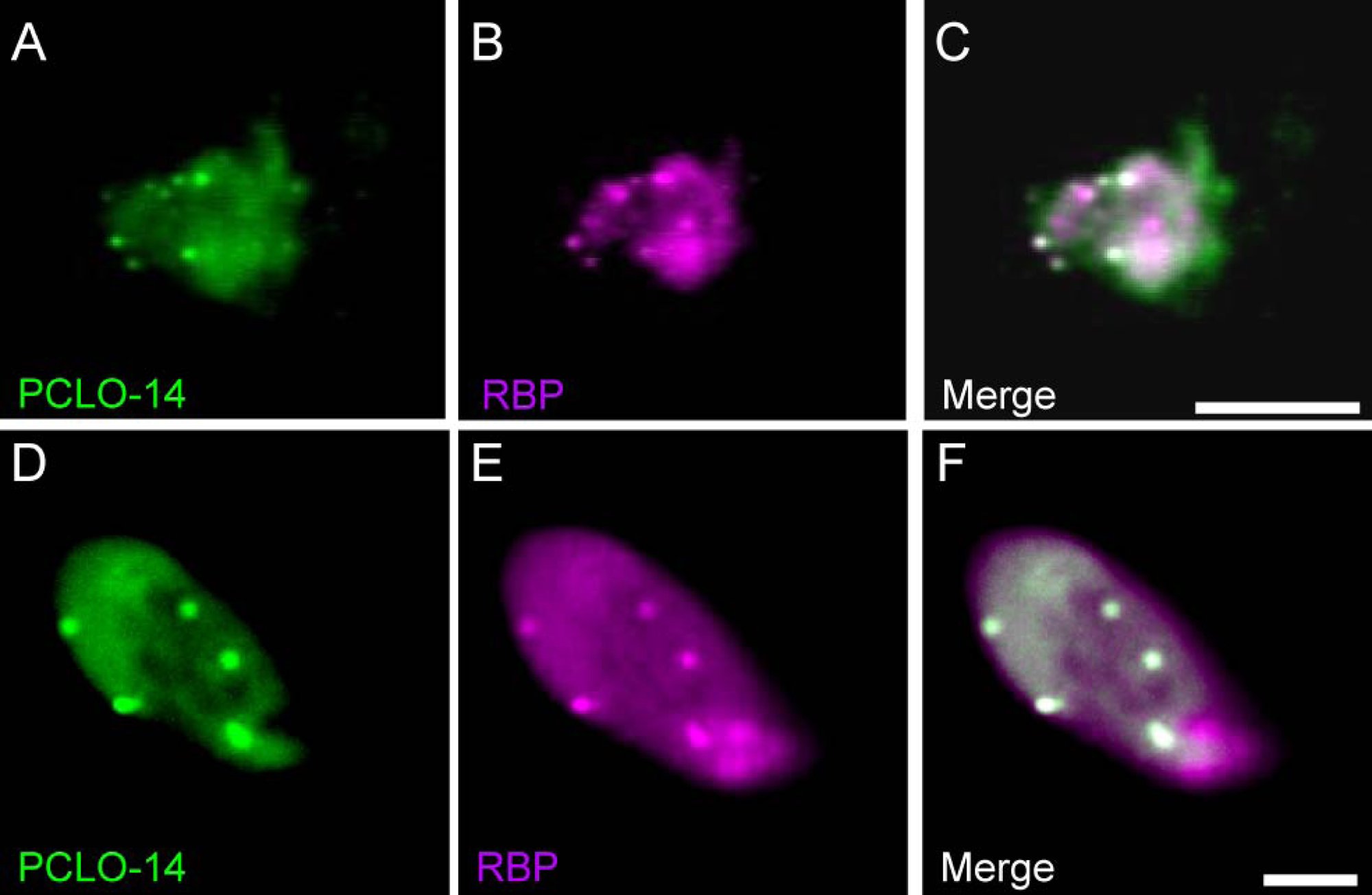
Piccolo PVDLT containing peptide labels ribbons in bipolar cells. Goldfish retinal bipolar cells were loaded with a fluorescent Ribbon-binding peptide (RBP; magenta) and a 14 amino acid peptide derived from Piccolo that encompasses the PVDLT motif (PCLO-14; green) and imaged using TIRF microscopy. (A) bright-field image of bipolar cell terminal. (B) Merged TIRF image of PCLO-14 (green) and RBP (magenta) peptides. Same cell as in A, C, D. Scale bar: 10 µm in B.

### PXDLS/T-binding mutants fail to localize to synaptic locations

We next sought to study the effects of PXDLS/T-binding mutants on ribbons in vivo. To do so, we turned to a zebrafish neuromast hair cell preparation that is amenable to rapid and easy transgene expression and electrophysiology (Ricci et al., 2013; Sheets et al., 2017; Kindt and Sheets, 2018). We introduced WT and mutant versions of the short isoform of Ribeye with a YFP tag (Ribeye(a)s-YFP) into zebrafish mutants harboring homozygous frame-shift mutations in both *ribeye* genes (*ribeye [a(*Δ*10)/b(*Δ*7)];* Lv et al., 2016) and marked post-synaptic locations using a pan-MAGUK antibody (Meyer et al., 2005; Sheets et al., 2011).

In Ribeye(a)s-YFP overexpressing (*ribeye [a(*Δ*10)/b(*Δ*7)]*, pan-MAGUK staining was found near to bright spots immuno-positive for Ribeye (figure 6). By contrast, fluorescent spots in Ribeye aV338R-YFP exhibited less apparent co-localization (Figure 6B). To quantify, we measured the distance between the center of individual Ribeye spots to the nearest postsynaptic pan-MAGUK spot (Figure 6G). Based on this analysis, we found that Ribeye(a)s-YFP spots were on average 0.22±0.03µm (n = 20) from their nearest pan-MAGUG spots, whereas Ribeye a V338R spots were 0.96±0.20 µm (n=21)..

**Figure 6.**
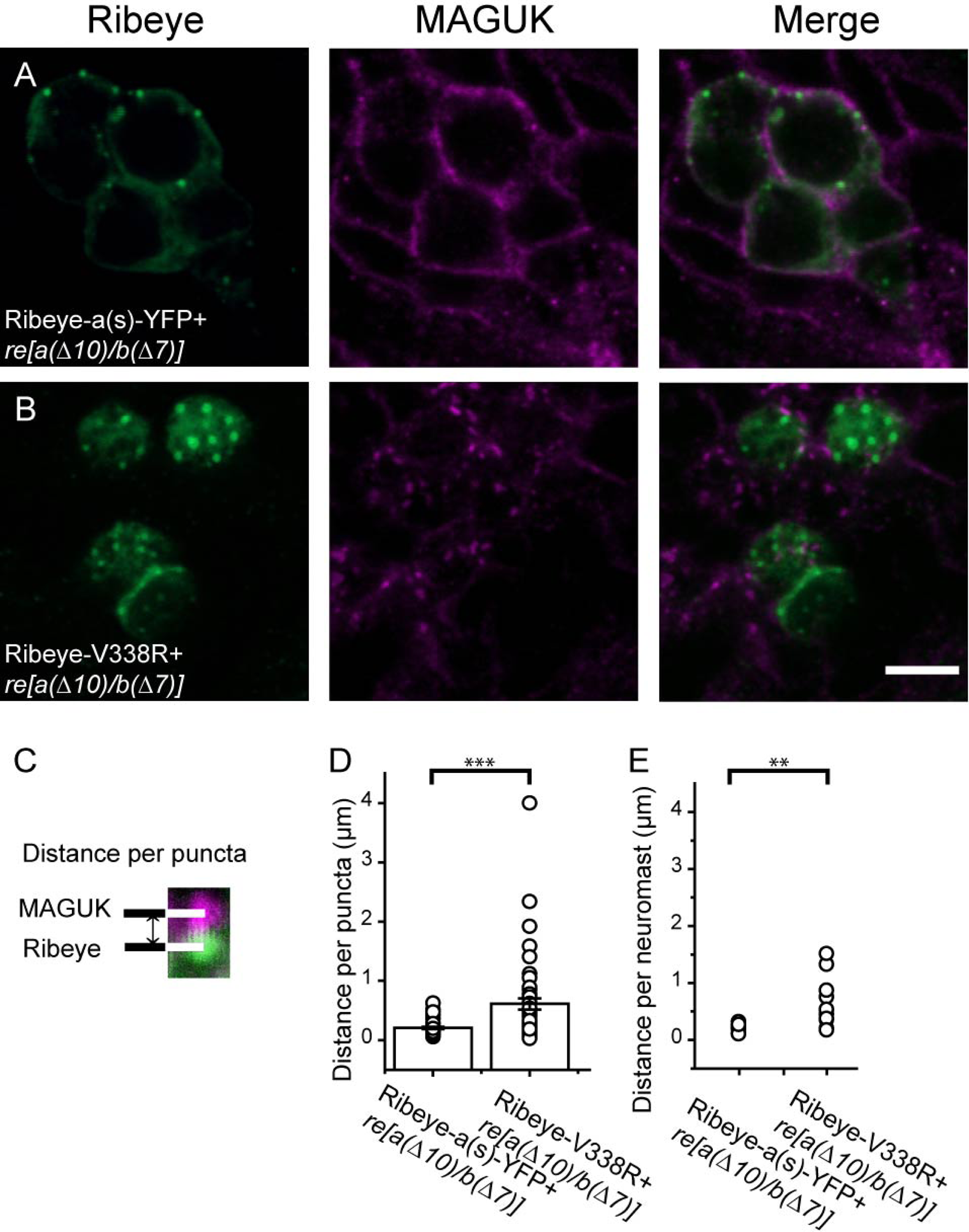
Ribeye-Piccolo colocalization requires intact PXDLS/T-binding cleft. A-B: *ribeye (a(Δ10)/b(Δ7)* zebrafish were injected with plasmids for either the short isoform of Ribeye a concatenated to YFP (Ribeye as-YFP) or to the short isoform of Ribeye a with a mutation to its PXDLS/T-binding cleft (Ribeye as(V338R)-YFP) co-stained with the post-synaptic marker pan-MAGUK. (A) Expression of WT-Ribeye in the double-homozygous mutant zebrafish detected by ctbp2 antibody shows a typical colocalization to postsynaptic density marker pan-MAGUK. (C) and (D) Enlargement of ROIs shown in A; (E) and (F) magnified image of ROI in (B). (G) To determine synaptic localization of Ribeye spots, the distance of each Ribeye spot was measured to the nearest pan-MAGUK spot. (H) Circles denote distances between each individual Ribeye spot and nearest pan-MAGUK spot, bar graph represents the mean across spots. (I) same as in H, but showing distributions of averages across neuromasts. Error bars indicate SEM. ***P<0.001, **P<0.01.

### Morphology of Hair Cell Synaptic Ribbons with PXDLS/T-binding Mutants

We next looked at the ultrastructure of hair cells overexpressing wild type and mutant Ribeye in the (*ribeye [a(*Δ*10)/b(*Δ*7)]* double homozygous mutants. Ribbons in neuromast hair cells in 5 dpf zebrafish are spherical and localize near the plasma membrane opposite afferent fibers (Obholzer et al., 2008). As expected, we observed typical dense core ribbons EM sections of neuromasts hair cells in control WT fish. By contrast, double homozygous mutant *ribeye [a(*Δ*10)/b(*Δ*7)]* maintained an organized collection of synaptic vesicles in a spherical array, but lack the electron density (Figures 7 AB), as previously described (Lv et al., 2016). These structures, referred to as ‘ghost-ribbons’, are smaller than WT ribbons and are often mislocalized away from afferents and sometimes appear as clusters rather than well-separated individual units [(Lv et al., 2016) and see figure 8F].

**Figure 7.**
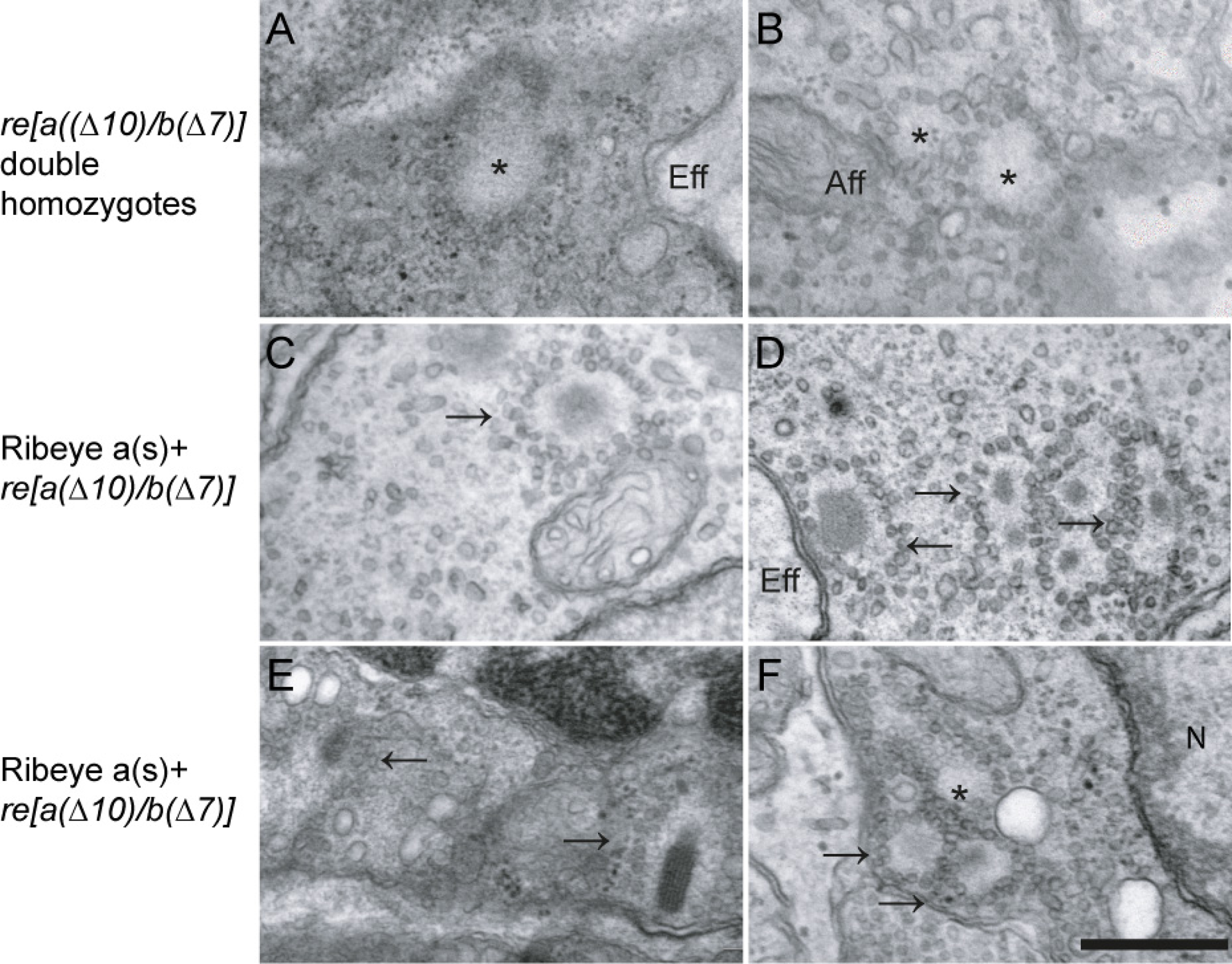
Ribeye(a)s-YFP overexpression partially rescues the lost synaptic density in ribbons of neuromast hair cells in *ribeye(a(Δ10)/b(Δ7)* mutants. (A-B) Electron micrograph showing typical examples of ghost ribbon (denoted by *) in neuromast hair cells in *ribeye(a(Δ10)/b(Δ7))* zebrafish mutants. (C-F) Examples of electron micrographs from Ribeye(a)s-YFP over-expressing hair cells in the in *ribeye(a(Δ10)/b(Δ7))* zebrafish mutants. Note the electron densities in the structures with diverse morphologies (arrows). Most densities were spherical, but were smaller than densities in WT ribbons, but some showed bar-like laminar structures (e.g. lower right density in E), reminiscent of retinal ribbons. All images taken from 5 days post-fertilization fish. Scale bar is 500 nm. Note that despite partial return of density to ribbons, many ribbons still fail to localize to synaptic locations.

**Figure 8.**
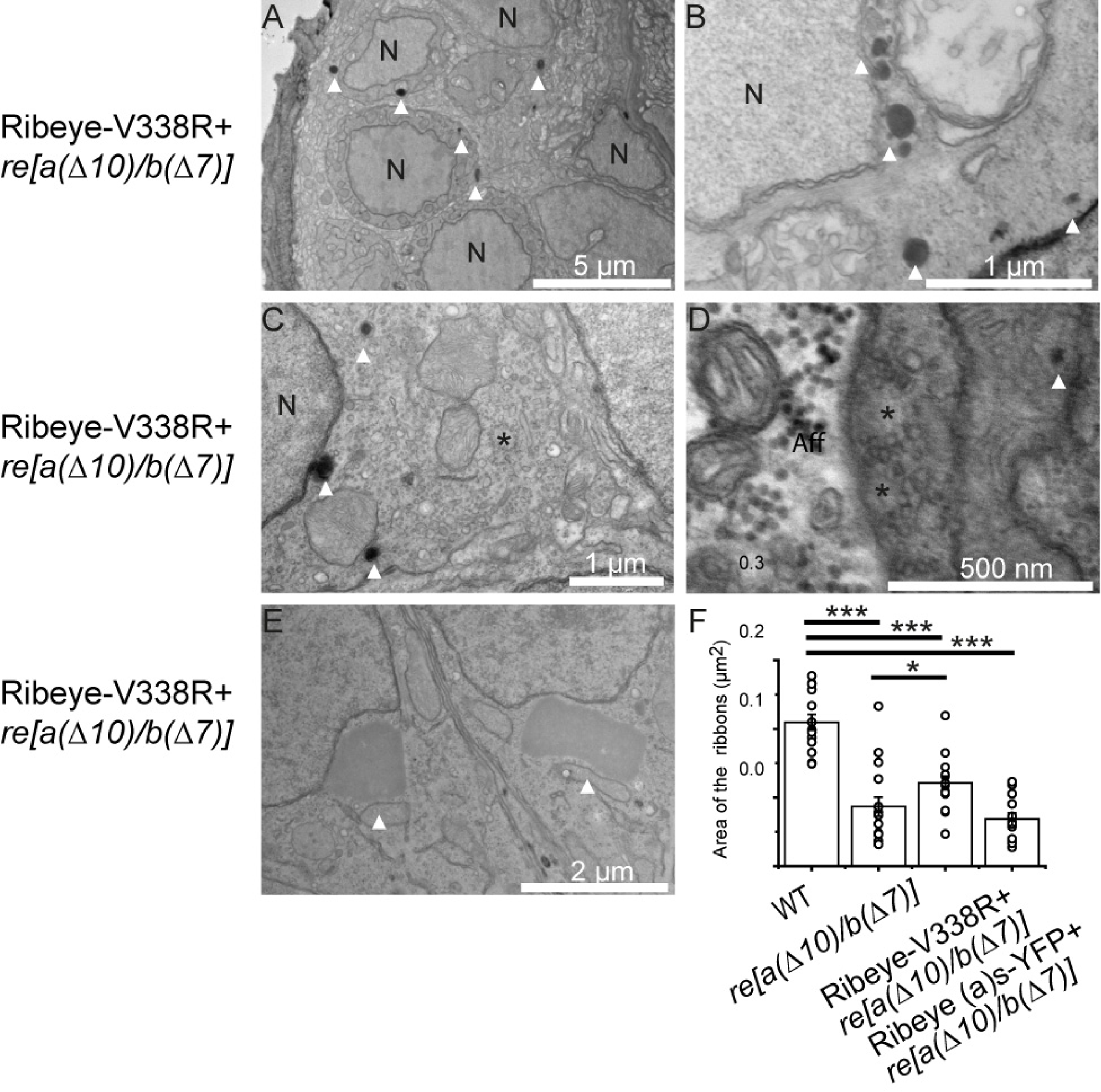
Ribeye as-(V338R)-YFP overexpression fails to rescue the lost synaptic density in ribbons of neuromast hair cells in *ribeye(a(Δ10)/b(Δ7)* mutants. A-E: Examples of electron micrographs from WT Ribeye as-(V338R)-YFP over-expressing hair cells in the in *ribeye(a(Δ10)/b(Δ7))* zebrafish mutants. A) low magnification image showing that electron densities scattered throughout neuromast of Ribeye as-(V338R)-YFP expressing hair cells. (B, C) Higher magnification images showing that unlike Ribeye(a)s-YFP expressing cells (figure 7), electron densities are not surrounded by vesicle-containing ribbon-ghosts. (D) Example of ribbon-ghosts lacking a density (denoted by *) in Ribeye as-(V338R)-YFP expressing cell. (E) Example of giant electron dense structures occasionally found in Ribeye (a)s-(V338R)-YFP expressing cells. (F) Areas of ribbons or ghost-ribbons in WT, *ribeye(a(Δ10)/b(Δ7))* and *ribeye(a(Δ10)/b(Δ7))* overexpressing either Ribeye(a)s-YFP or Ribeye (a)s-(V338R)-YFP. Note that Ribeye (a)s-YFP causes an increase in ribbon size, but ribbons still remain smaller than WT ribbons. *P<0.05, **P<0.01, ***P<0.001.

Electron micrographs of *ribeye [a(*Δ*10)/b(*Δ*7)]* fish overexpressing WT Ribeye A-short exhibited electron densities in the cores of ribbon ghosts. The size and shape of these electron densities varied (Figure 7C-F), with most showing round densities within the core (Figure 7C-D), but with some showing more rectangular and laminar structures, somewhat reminiscent of retinal ribbons (Figure 7E). In general, Ribeye(a)s-YFP overexpression in the *ribeye [a(*Δ*10)/b(*Δ*7)]* mutants led to smaller densities than those observed in WT animals (Figure 9) and many micrographs showed aberrant clusters and mislocalized ribbons (Figure 7D&F). We found 14 ghost ribbons with electron densities and no obvious examples of ectopic electron dense spots in these fish. Ghost ribbons containing dense cores (0.12 ±0.01µm^2^ n=18) were significantly larger than ghost ribbons in *ribeye [a(*Δ*10)/b(*Δ*7)]* double homozygous mutants(0.09±0.01 µm^2^ n=16) that were not expressing Ribeye (Figure 8F), but still smaller than the densities in WT ribbons(0.21±0.01 µm^2^ n=14). Overall, our results suggest that Ribeye(a)s-YFP in the *ribeye [a(*Δ*10)/b(*Δ*7)]* mutant lines resulted in partial recovery of ribbon morphology, which may reflect either the need for additional or different isoforms of Ribeye or higher expression levels of Ribeye.

**Figure 9.**
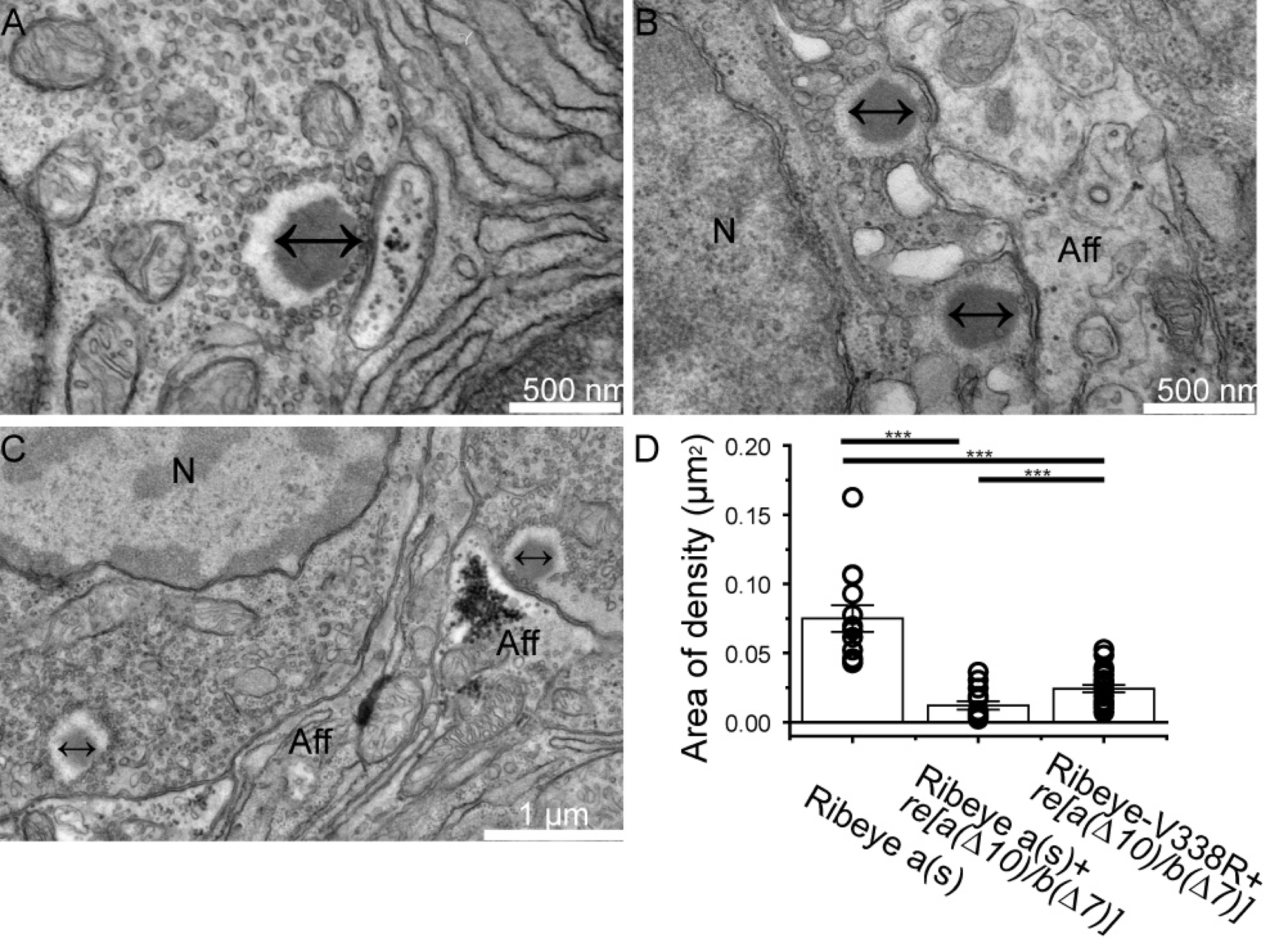
Measurements of electron densities. (A-C) Examples of electron micrographs of WT ribbons in hair cells of zebrafish neuromasts. (D) Area of electron densities in WT animals and in *ribeye(a(Δ10)/b(Δ7))* mutants overexpressing either Ribeye(a)s-YFP or Ribeye(a)s-(V338R)-YFP. Note that electron densities in Ribeye(a)s-(V338R)-YFP were outside of ribbon structures.

In contrast to Ribeye(a)s-YFP expressing fish, those overexpressing the Ribeye A short V338R-YFP in *ribeye [a(*Δ*10)/b(*Δ*7)]* double homozygous fish exhibited no densities in their ghost ribbons(Figure 8D), but instead we found 26 ectopic electron densities that lacked synaptic vesicles in 8 electron micrographs (Figure 8). Most appeared as small electron-dense structures, but a subset (figure 8A-C) were nearly 2 μm and were reminiscent of ectopic structures previously reported in neuromast hair cells that overexpress high levels of Ribeye. These results suggest that Ribeye a(s) V338R-YFP is unable to be recruited to ghost ribbons (Figure 8E; (Sheets et al., 2011), perhaps due to its inability to bind to Piccolino.

### PXDLS/T mutation affects the release properties of ribbon synapses

We next tested whether the PXDLS/T-binding mutants cause dysfunction in synaptic release properties. To study Ribeye and its PXDLS/T binding cleft function, we generated mosaic zebrafish overexpressing WT-Ribeye a short (Ribeye(a)s-YFP) or mutant Ribeye (Ribeye a(s) V338R-YFP) in double homozygous *ribeye [a(*Δ*10)/b(*Δ*7)]* fish. Hair cells containing YFP were targeted for recording and non-expressing cells lacking YFP in the same fish were used as controls. Whole-cell membrane capacitance recordings were performed in hair cells of these fish at 4-9 dpf.

The average of capacitance response from the hair cells of *ribeye [a(*Δ*10)/b(*Δ*7)]* fish without fluorescence (n=20) showed a capacitance increase of 80 ± 4.5 fF. in response to a 3-second step depolarization to −10mV (Figure 10), Ribeye a(s)-YFP expressing cells exhibited a smaller response. The peak amplitude of the capacitance of Ribeye a(s)-YFP is 30 ± 3.5 fF, n=8, significantly smaller than non-fluorescent cells (p=.0001). WT zebrafish show a similar recorded capacitance to the Ribeye a(s)-YFP overexpressing cells in *ribeye [a(*Δ*10)/b(*Δ*7)]* mutants (30.1±4.3Ff,n=5). By contrast YFP positive cells in those injected with the Ribeye a V338R-YFP constructs averaged an increase of 62.6 +/− 8.7 fF, not significantly different than non-fluorescent cells (p = 0.3)

**Figure 10.**
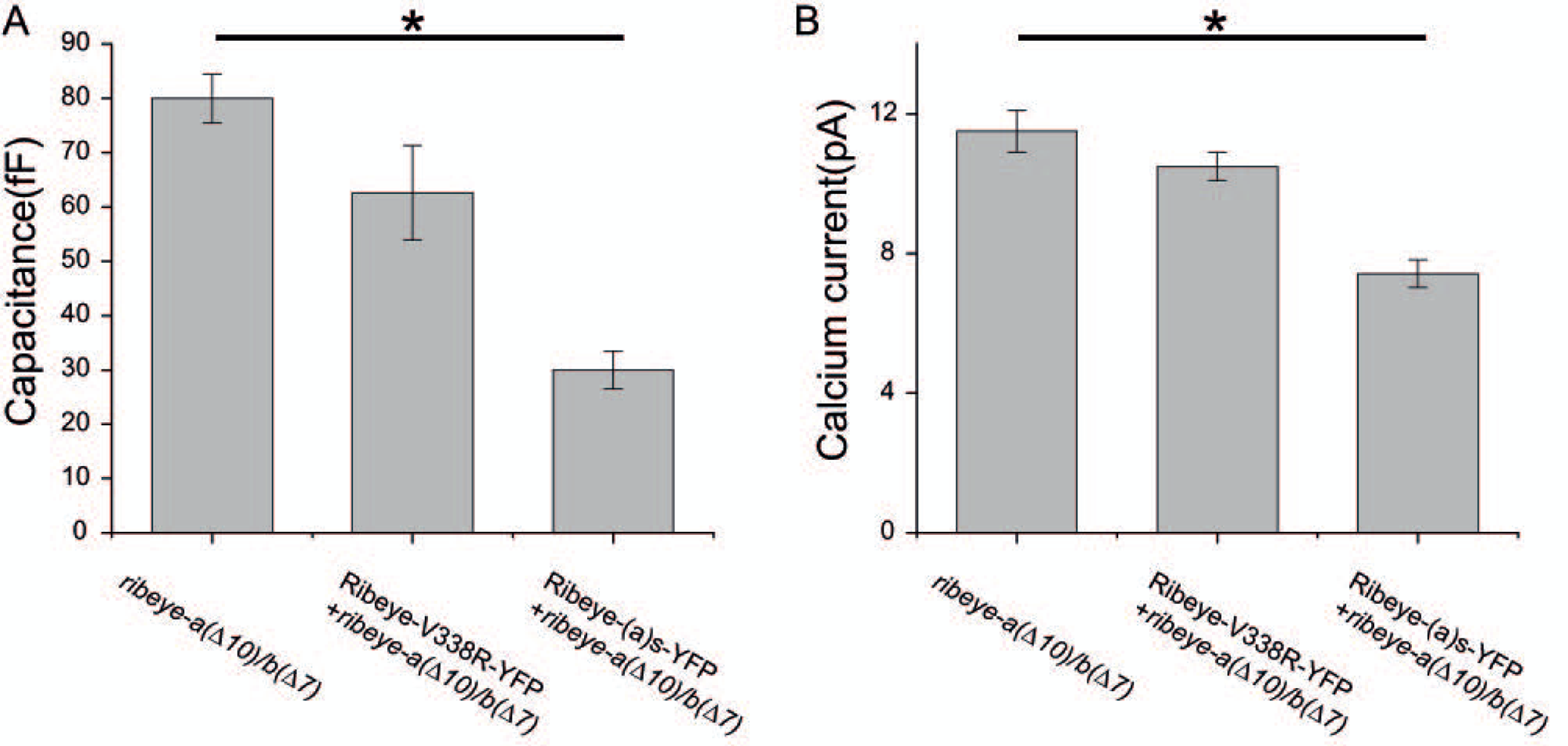
Lateral line hair cells overexpressing Ribeye(a)s-YFP, but not those expressing Ribeye(a)s-(V338R)-YFP reduce calcium currents and capacitance responses. Neuromast hair cells from 4-9 dpf zebrafish were whole-cell voltage clamped and capacitance measurements were performed to track exocytosis. Hair cells were subjected to 3 s long depolarization to −10 mV and the difference after the depolarization relative to the baseline before stimulation was measured. Zebrafish embryos were injected at the one cell stage with a construct encoding for either Ribeye-as-YFP or Ribeye(a)s-(V338R)-YFP. Fluorescent cells containing the transgenes were compared against non-fluorescent cells in the same fish as controls. (A) A shows the average capacitance recordings of hair cells expressing Ribeye-as-YFP and Ribeye(a)s (V338R)-YFP in zebrafish(a(Δ10)/b(Δ7). (B) Hair cells were depolarized to −20 mV and leak-subtracted currents were measured for same animals as in A. Bar graphs show mean +/− SEM. *P<0.05.

In our previous study, we found that *ribeye [a(*Δ*10)/b(*Δ*7)]* had a slightly larger calcium current than WT fish at a membrane potential of −20 mV (Lv et al., 2016). We tested whether either WT Ribeye a(s)-YFP or Ribeye a(s) V338R-YFP could reduce calcium currents.. The average current in response to voltage steps to −20 mV was 11.5 ± 0.6 pA (n = 17) in non-fluorescent hair cells. The calcium current was smaller in Ribeye(a)s-YFP expressing cells 7.42 ± 0.4 (n = 8), with the difference in means reaching a marginal level of significance (p = 0.047). By contrast, Ribeye as V338R-YFP expressing cells showed calcium currents that were much closer to non-expressing cells 10.9 ± 0.4 pA (n = 10; p = 0.77) (Figure 10).

## Discussion

Ribeye is a major component of the synaptic ribbon (Schmitz et al., 2000), making up the majority of the protein in the structure (Zenisek et al., 2003). Ribeye is transcribed using an alternative start site of the transcriptional co-repressor, CtBP2 and retains a B-domain that is identical to CtBP2, except for lacking CtBP2’s N-terminal 20 amino acids (Schmitz et al., 2000). Disruption of Ribeye leads to dramatic effects on synaptic ribbon morphology in zebrafish and mouse (Wan et al., 2005; Sheets et al., 2011) (Lv et al., 2016; Becker et al., 2018; Jean et al., 2018; Mesnard et al., 2022) and proper localization of calcium channels and the presynaptic protein RIM2 to synaptic locations (Sheets et al., 2011; Lv et al., 2012; Maxeiner et al., 2016; Jean et al., 2018). Surprisingly genetic disruption of Ribeye is accompanied by comparatively subtle effects on exocytosis in hair cells (Lv et al., 2016; Becker et al., 2018; Jean et al., 2018) and a reduction in amplitude, but not kinetics of EPSCs evoked by release from retinal bipolar cells (Maxeiner et al., 2016). Interestingly, the ribbon has long been suggested to facilitate prolonged exocytosis, the maximal rates of exocytosis in hair cells of both fish and mouse are either unaffected or slightly augmented when depleted of Ribeye. Reduced rates of exocytosis are only apparent in these cells for more modest stimuli, suggesting a rethinking of the role of the ribbon in exocytosis. Despite the mild effect on exocytosis, mice exhibit a mild hearing loss, a reduction in firing rate and a slower recovery from adaptation in the threshold for hearing and changes in the frequency response in Ganglion cells in the retina (Becker et al., 2018; Jean et al., 2018; Okawa et al., 2019). The emerging results suggest an important role for the ribbon in sensory processing different from the assumed role of a simple facilitator of long-term release. Hence, the role of Ribeye specifically and the ribbon, more generally, remains enigmatic.

The CtBP (C-terminal binding protein) family of proteins was originally named due to the founding member’s (CtBP1) interaction with the C-terminus of adenoviral E1A proteins. The interaction site was determined to require a PLDLS motif that is highly conserved in E1A proteins across many adenoviruses (Schaeper et al., 1995). Subsequently, it has been found that both CtBP1 and CtBP2 interact with not only viral proteins, but numerous other proteins harboring this sequence and these interactions are critical for many CtBP functions in the nucleus (Quinlan et al., 2006; Zhao et al., 2007; Kuppuswamy et al., 2008). Crystal structures of CtBP1 with PXDLS/T peptides reveal that these peptides interact in a specific binding cleft on the surfaces of the CtBPs (Nardini et al., 2006). Mutations to residues of this cleft prevent interactions for CtBPs to bind PXDLS/T containing partner proteins (Zhang et al., 2000; Molloy et al., 2007). As noted above and in previous work (Muller et al., 2019), the synaptic ribbon protein Piccolino contains a well conserved PVDLT sequence and several PXDLS-like motifs, suggesting an interaction between Ribeye and Piccolino via an analogous interaction. Indeed, Muller et al. (2019) showed using several techniques an interaction between the two proteins using these sites. Here, we provide additional evidence of this interaction and investigate whether the PXDLS/T-binding cleft is necessary for normal Ribeye function (Muller et al., 2019).

We show here that Ribeye, but not a PXDLS/T-binding cleft mutant, colocalized with Piccolo in INS-1 cells and with Piccolo fragments co-transfected in HEK cells, similar to recently published results (Muller et al., 2019). Moreover, we found that a short peptide fragment localizes to ribbons in retinal bipolar cells when introduced into the cell via a patch pipette. Muller et al. (2019) also demonstrated using co-IP and co-expression studies that in addition to the PVDLT domain numerous similar motifs that varied by single amino acids from the canonical motif needed to be mutated to abrogate Ribeye-Piccolino interactions. Together these results are strong evidence for a direct interaction between these two proteins.

Our previous work demonstrated that introduction of frame-shift mutations into both zebrafish *ribeye* genes *(ribeye [a(*Δ*10)/b(*Δ*7)])* led to loss of most Ribeye protein in hair cells and loss of the electron dense core in synaptic ribbons, mislocalization of ribbons and a modest enhancement in synaptic exocytosis (Lv et al., 2016). We used these mutants as a mostly ‘blank slate’ to introduce both wild-type and mutant Ribeye to explore whether mutant Ribeye can properly target to synaptic ribbons or rescue normal function. We deemed it necessary to use these mutants, since previous work identified numerous interaction sites between the A-domain and B-domain of Ribeye (Magupalli et al., 2008) and a previous study demonstrated that the B-domain and some A-domain isoforms localize to synapses in zebrafish hair cells (Sheets et al., 2014). It is worth noting that at least 3 isoforms of Ribeye have been identified in zebrafish: two splice variants of the *ribeye a* gene and two variants of the *ribeye b* gene (Wan et al., 2005). Additionally, a total of 16 splice variants of the *CtBP2* gene, including 6 variants containing portions of the ribbon-specific A-domain are predicted based on genomic sequencing (www.ensembl.org).

We find here that overexpression of Ribeye as-YFP in the *ribeye [a(*Δ*10)/b(*Δ*7)]* double homozygous line leads to a return of electron densities to ribbon synapses and normal levels of exocytosis and calcium currents. The rescue appears incomplete, as the ‘rescued’ ribbons exhibited smaller densities, some unusual morphologies (figure 7D, E) and imprecise localization. We hypothesize that the incomplete rescue is due to either 1) a reduction in overall Ribeye expression level compared to wild type animals or 2) the requirement for multiple isoforms of Ribeye A and/or Ribeye B to fully restore the ribbon.

In contrast to the Ribeye as-YFP expressing cells, Ribeye as (V338R)-YFP expression did not lead to the restoration of electron densities to ribbons nor normal exocytosis or calcium currents in the *ribeye [a(*Δ*10)/b(*Δ*7)]* mutants. Moreover, ectopic electron dense aggregates without synaptic vesicles were found in these overexpressing cells. These structures are similar to ectopic densities seen previously in zebrafish hair cells highly overexpressing Ribeye (Sheets et al., 2011; Sheets et al., 2017) and in cell lines (Schmitz et al., 2000; Magupalli et al., 2008) expressing Ribeye. To explain these results, we suggest a model that Piccolino recruits Ribeye to synaptic ribbons and in the absence of Piccolino (in the case of cell lines) or saturation of Ribeye interaction sites (in the case of hair cells overexpressing Ribeye), Ribeye forms cytoplasmic electron-dense aggregates away from the synapse. Interestingly, neither the aggregates formed by Ribeye as (V338R)-YFP overexpression in *ribeye [a(*Δ*10)/b(*Δ*7)]* mutants here nor the aggregates formed by Ribeye overexpression in WT animals (Sheets et al., 2011; Sheets et al., 2017) localized to synaptic regions. Relatedly, Ribeye-deficient ghost-ribbons exhibit some degree of synaptic localization, but are often mislocalized to cytoplasmic and non-synaptic membrane locations (Lv et al., 2016). These results suggest that Ribeye itself is insufficient for synaptic localization, but that adequate levels of Ribeye may be necessary to maintain or stabilize ribbons at synaptic locations and recruit other proteins to synaptic locations. Indeed, ectopic Ribeye aggregates have been shown to colocalize with calcium channels (Sheets et al., 2011) and genetic disruption of Ribeye has been shown to have effects on the coupling of calcium channels to exocytosis (Lv et al., 2016; Maxeiner et al., 2016) and calcium channel properties (Becker et al., 2018; Jean et al., 2018; Grabner and Moser, 2021). Further experiments will be necessary to test this model.

The interaction between Ribeye and Piccolo also have implications for ribbon structure. While it has been known for some time that Ribeye alone can form electron dense aggregates (Schmitz et al., 2000; Magupalli et al., 2008), Muller et al. (2019) recently showed that co-expression of Ribeye and Piccolino in HEK cells can lead to bar-shaped structures that appear photoreceptor ribbon-like at the level of light microscopy. Although the previous work did not examine the structures at the electron microscopy level, it suggests that Ribeye-Piccolino interactions are likely to play a prominent role in determining the shape of the ribbon. However, the diversity of ribbon shapes across cell types remains unexplained.

The A-domain of Ribeye and synaptic ribbons are found in the hagfish, *Eptatretus burgeri,* an early chordate, but not earlier in closely-related invertebrates, such as the tunicate, *Ciona intestinalis* indicating that Ribeye and ribbons evolved after this evolutionary split (Zenisek and Thoreson, 2024). Interestingly, hagfish photoreceptors have spherical synaptic bodies rather than the platelike ribbons seen in retinas of true vertebrates (Holmberg and Ohman, 1976) and also lack the necessary PXDLT sequences necessary for Piccolo to interact with Ribeye. Although merely a correlation, this is consistent with a role for Piccolino/Ribeye interactions in driving ribbon structure (Zenisek and Thoreson, 2024).

## Acknowledgements

The authors would like to thank SueAnn Mentone for her technical help and expertise with electron microscopy. This work was funded by a grant from the National Institute of Health (EY032396) and the Yale University Vision Core (EY026878).

